# The baboon as a statistician: Can non-human primates perform linear regression on a graph?

**DOI:** 10.1101/2024.06.16.599196

**Authors:** Lorenzo Ciccione, Thomas Dighiero Brecht, Nicolas Claidière, Joël Fagot, Stanislas Dehaene

## Abstract

Recent studies showed that humans, regardless of age, education, and culture, can extract the linear trend of a noisy graph. Here, we examined whether such skills for intuitive statistics are confined to humans or may also exist in non-human primates. We trained Guinea baboons (*Papio papio*) to associate arbitrary geometrical shapes with the increasing or decreasing trends of noiseless and noisy scatterplots, while varying the number of points, the noise level, and the regression slope. Many baboons successfully learned this conditional match-to-sample task for both noiseless and noisy plots. Crucially, for successful baboons, accuracy varied as a sigmoid function of the t-value of the regression, the same statistical index upon which humans also base their answers, even after controlling for other variables. These results are compatible with the hypothesis that the human perception of data graphics is based on the pre-emption and recycling of a phylogenetically older competence of the primate visual system for extracting the principal axes of visual displays.

## Introduction

Humans are known to perform remarkably well at tasks involving statistical evaluations of their environment, an ability with clear evolutionary advantages (Gigerenzer & Murray, 2015). They correctly estimate the probability of the occurrence of events drawn from given distributions (Nisbett & Krantz, 1983; Xu & Garcia, 2008); they can recognize and learn statistical regularities in both linguistic (Benjamin et al., 2023; Saffran et al., 1996, 1999) and non-linguistic stimuli (Yuille & Kersten, 2006); they can extract the average of many different features such as color, orientation or size from large datasets of items, a set of skills that has been called “ensemble perception” (Alvarez, 2011; Whitney & Yamanashi Leib, 2018).

Some of these intuitive statistical abilities have been found in non-human primates as well. For instance, chimpanzees can make probability judgments based on several proportional ratios (Eckert et al., 2018), long-tail macaques can infer simple heuristics (Placì et al., 2018), capuchin monkeys can make probabilistic inferences (Tecwyn et al., 2017), and baboons can learn spatial statistical contingencies (Goujon & Fagot, 2013). Statistical learning skills are also undisputed in many other animal species (for a review, see Santolin & Saffran, 2018). However, whether non-human primates can also perform an ensemble evaluation of a simultaneously presented dataset is unknown and, to date, yet to be proven in other animal species.

We decided to test such a skill in a population of Guinea baboons (*Papio papio*), an Old-World monkey. In order to do so, we turned our attention towards a recently developed binary task of trend judgment over scatterplots (Ciccione & Dehaene, 2021). In this task, on each trial, humans had to judge, as fast and as accurately as possible, whether the underlying trend of a noisy scatterplot comprising from 6 to 66 dots was increasing or decreasing – a task equivalent to that of a statistician deciding whether there is a significant ascending or descending trend in the data. Results showed that both adults and 6-year-old children, whether schooled or unschooled, could perform this task and that, crucially, they based their judgment on the t-value of the dataset, the very index that a statistician would compute to estimate the significance of the trend (Ciccione et al., 2023). The t-value is a summary of several data features: it combines the signed slope (either positive or negative), the level of noise in the dataset (with noisier scatterplots resulting in lower t-values), and the number of points in the graph (the larger the number, the higher the t-value). The following formula provides the computation of the t value:

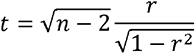

where n is the number of data points, and r indicates the correlation coefficient, i.e., the covariance of x and y values, divided by the product of their standard deviations (i.e., their respective noise levels).

In other words, the decision variable that humans use to evaluate the statistical trend of a noisy ensemble is exactly the one that a statistician would use (Peterson & Beach, 1967): it takes into account all of the relevant scatterplot features in their judgments, as if they were implicitly performing a t-test over the graph. Their estimate of the slope, however, is not unbiased: humans do not extract the exact regression line predicted by classical least squares regression, which minimizes the vertical distance of the points to the fit, but another line that minimizes the orthogonal distance of the points to the fit (Deming regression; Ciccione & Dehaene, 2021) and is mathematically equivalent to the calculation of the principal axis of the graph. During object perception, the principal axis is easily and automatically computed by both human (Ayzenberg et al., 2019; Bodily et al., 2018; Lowet et al., 2018) and non-human primates (Hung et al., 2012).

Taking these results together, it has been hypothesized that human intuitive graph perception skills might be based on a neuronal recycling of cortical areas devoted to the extraction of the orientation of the principal axis of objects (Ciccione & Dehaene, 2021). According to the neuronal recycling hypothesis (Dehaene & Cohen, 2007), cognitive processes and brain areas initially devoted to a specific computation may be reused for a novel cultural task, providing that the novel function is sufficiently close to the initial one. For example, circuits for approximate number sense seems to be repurposed for human symbolic mathematics, and circuits for the fast visual recognition of objects for reading (Amalric & Dehaene, 2016; Dehaene, 2005; Hannagan et al., 2021; Rajalingham et al., 2020).

Similarly, here, we test the idea that the extraction of object orientation underlies the capacity to perceive the linear trend in a data graph. While non-human primates are certainly not exposed to such graphs in their daily lives, they are able to orient their hands in order to grasp objects, which implies an extraction of their principal axis. We propose that this capacity could be extended to trend judgments over noisy graphs, provided that they consider the set of dots as a single “object”.

With those ideas in mind, here we tested whether non-human primates (baboons) could perform an intuitive statistical trend judgement over noisy ensembles. We thus taught to a group of 23 Guinea baboons to associate, in a series of conditional match-to-sample tasks an arbitrary stimulus to ascending trends and another arbitrary one to descending trends. Such trends could be either noiseless (thus represented by straight lines) or hidden in noise. On each trial, the two response stimuli among which the animal had to choose were presented randomly on the left or on the right of the screen, thus preventing baboons from systematically using only a subset of the dots and choosing the response stimulus that was closer to them. To anticipate on the results, we found that baboons could learn such an association for both noiseless and noisy trends, and, crucially, that they based their judgment on the t-value, the same statistical index used by humans to perform the task.

## Methods

### Participants

A group of 23 *Papio papio* baboons, comprising 16 females with ages ranging from 89 to 309 months and a mean age of 174 months (14.5 years), underwent testing at Comparative Cognition platform of the “Centre de Recherche en Psychologie et Neuroscience” situated at the CNRS “Station de Primatologie” facility in Rousset-sur-Arc, France. The baboons inhabited a spacious 700-square-meter outdoor enclosure, complemented by access to indoor housing. Additionally, they had the opportunity, on a voluntary basis, to engage with 14 automated learning devices, located inside small covert cabins (Fagot & Bonté, 2010; Fagot & Paleressompoulle, 2009). These devices featured a 19-inch touch screen, a food dispenser, and a radio-frequency identification (RFID) reader capable of identifying individual animals. This identification allowed to keep track of each animal’s progression in the experimental sessions and thus, for each new entrance in one of the cabins, to make sure each baboon restarted from where they left. Overall, such experimental setup allowed for a versatile and enriched environment, encouraging the baboons to interact with the learning devices at their own discretion. This study received approval from the French Ministry of National Education (approval no. APAFIS-2717-2015111708173794-157 V3). Data from all participants (and the scripts for data analysis) are available on the OSF platform (https://osf.io/dyc6v/).

### Experimental procedure

The test was divided into three successive phases: a training phase followed by two experiments, each subdivided into a noiseless phase and a noisy phase (see Figure 1). Each monkey had to pass a criterion of at least 80% of correct responses in a given block of 136 trials in order to pass to the following experimental phase. The two experiments differed in terms of difficulty: the first one used the exact same stimuli previously used with human subjects (Ciccione et al., 2023), while the second, performed later, comprised stimuli with larger slopes and smaller levels of noise, as explained below.

**Figure 1.**
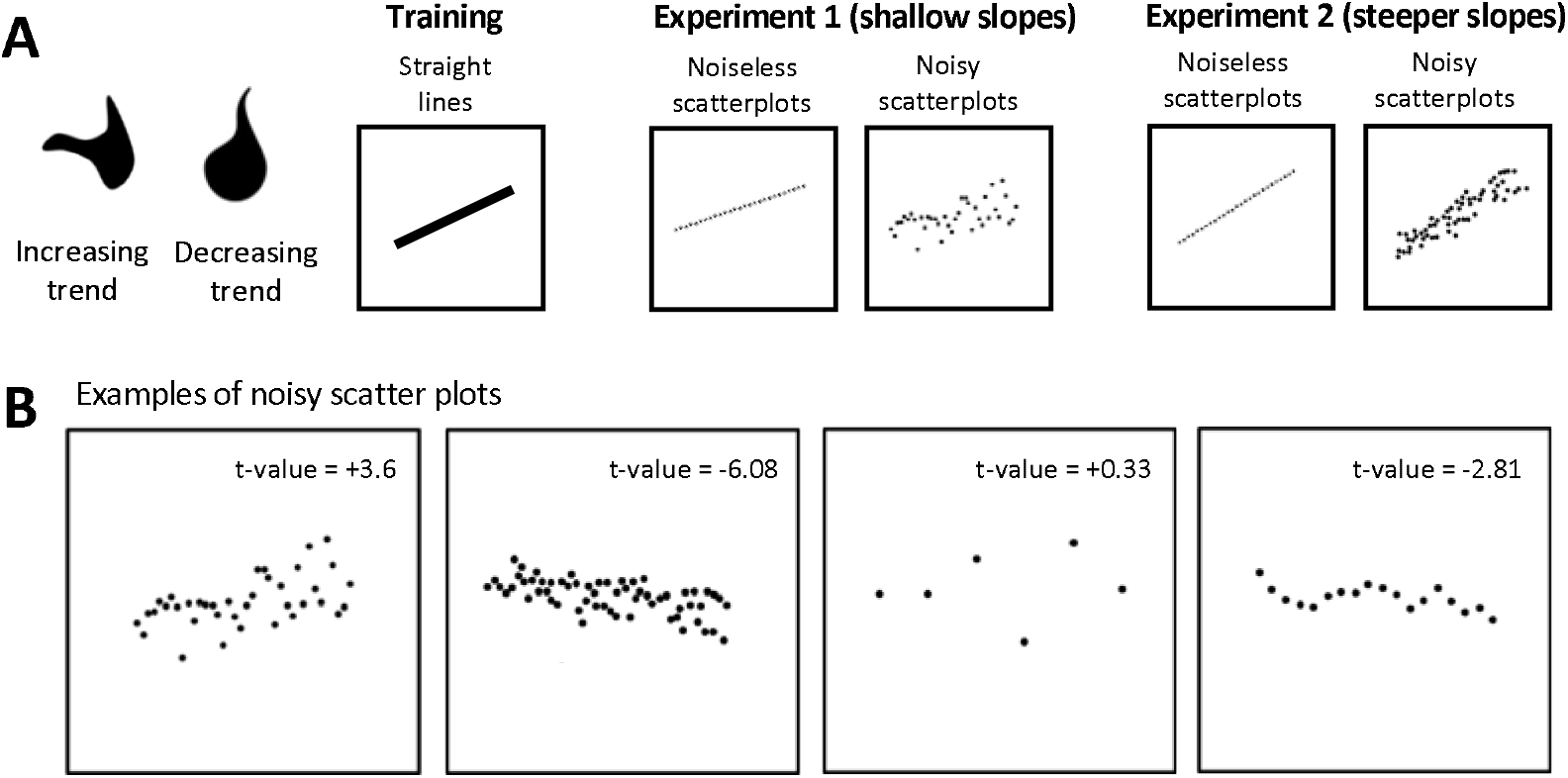
Experimental procedure to test linear trend perception from graphics in non-human primates. **A, training and testing phases.** On each trial, a graphic appeared and, after touching the screen, the animal had to choose between two shapes, one of which was arbitrarily associated with increasing trends and the other with decreasing trends. In the training phase, only straight lines oriented at -30° or +30° were presented. In experiment 1, scatterplots with shallow slopes matching those used in human experiments (Ciccione et al., 2023), were presented, either noiseless or noisy. In experiment 2, steeper scatterplots were presented, again either noiseless or noisy. **B: Examples of noisy scatterplots.** The number of dots, slope, and noise level could vary. In each example, we provide the associated t-value for illustration. Stimuli are plotted black on a white background for readability purposes but they were white on a black background in the experiment.

### Training

In this phase, baboons were familiarized with the conditional match-to-sample task that was used all along the different sessions. At the beginning of the trial, the baboons had to touch a fixation cross displayed at the bottom of the screen. Once done, it was presented with a straight line (white on black background) on the upper part of the screen, either oriented upwards at +30° or downwards at -30°. Touching the line produced the display of a third screen on which the two response stimuli were shown. One was the expected correct answer for lines oriented upwards, the other one was the expected correct answer for lines oriented downwards. The pair of response stimuli were randomly selected for each baboon among a list of 10 possible geometrical shapes. For a given baboon, the same pair was used for the entire duration of the experiment, but each stimulus could randomly appear either at the left or at the right of the screen, in order to avoid preferential response sides. The response stimuli we generated were inspired by the lexigrams taught to Kanzi, the famous bonobo (Savage-rumbaugh, 1996) (see Figure S3). A reward consisting of a small amount of food was given to the baboons if they provided a correct answer. Incorrect responses produced a 3 sec time-out indicated by a green screen. When a threshold of 80% of correct responses in a given block of 136 trials was reached, the first experiment started.

### Experiment 1, shallow slopes, noiseless scatterplots

During this phase, baboons were presented with straight lines made of 27 dots and with slopes varying from -18.75° to +18.75° (specifically: -18.75°, -12.5°, -6.25°, +6.25°, +12.5°, and +18.75°). Again, when subjects reached a criterion of 80% correct responses over a block of trials, they could pass to the following phase.

### Experiment 1, shallow slopes, noisy scatterplots

In this phase, baboons were presented with noisy scatterplots, again with the task of detecting whether the underlying trend was increasing or decreasing, by pressing one of the two response stimuli learnt during the training phase. The scatterplots measured 18.8 cm X 15.05 cm maximum on the screen. Given the viewing distance of 27.5 cm, their visual size was of 34.4° on the horizontal axis, and 28.6° on the vertical axis. The stimulus generation algorithm was exactly the same as the one previously used with human participants (Ciccione et al., 2023). Each scatterplot represented a dataset generated randomly from a linear equation (y_i_ = α x_i_ + ε_i_), where α denoted the prescribed slope, and ε_i_ were random numbers drawn from a normal distribution centered on zero with standard deviation σ. In a block, participants were presented with a total of 136 scatterplots resulting from combinations of three orthogonal factors: 7 prescribed slopes (α = − 0.1875, − 0.125, − 0.0625, 0, + 0.0625, + 0.125, or + 0.1875); 5 noise levels (σ = 0, 0.05, 0.1, 0.15, or 0.2); and 4 numbers of points (n = 6, 18, 38, 66). The 4 combinations with both a slope and a noise level of 0 were not presented (because the correct response would have been undefined). The x-axis coordinates were uniformly spaced and fixed for each n level. Figure 1B provides illustrative examples of stimuli, with the t-value for the Pearson coefficient of correlation indicated next to each scatterplot. This phase was prematurely interrupted after two months due to a too high difficulty of the task (most baboons had a score lower than 55%, although significantly different from 50%).

### Experiment 2, steeper slopes, noiseless scatterplots

All baboons, independently from their performance, were re-exposed to an easier version of the first phase of experiment 1, differing in the slope values that were twice as large: -37.5°, - 25°, -12.5°, +12.5°, +25°, +37.5°. For experiment 2 the size of the stimuli on screen was also slightly reduced (15 x 12 cm).

### Experiment 2, steeper slopes, noisy scatterplots

This testing phase was, again, identical to the previous corresponding phase except for the parameters used to generate the stimuli: to make the task easier, the slope values were doubled (-0.375, -0.25, -0.125, 0, +0.125, +0.25, +0.375) and the noise levels were halved (0, 0.025, 0.05, 0.075, 0.1). The number of points was the same (6, 18, 38, 66).

## Results

### Baboons learn to associate arbitrary shapes to ascending and descending straight lines

We first looked at the training phase, in which baboons had to learn to associate arbitrary shapes to the orientation of continuous lines with two possible slopes (-30° and +30°), and at the two phases that used noiseless scatterplots in which stimuli were a straight line of 27 dots with 6 possible slopes (experiment 1: -18.75° to +18.75°; experiment 2: -37.5° to +37.5°). As seen in figure 2 (left panel), performance in the training phase increased across trials. Overall, the 23 baboons had an above-chance accuracy of 59% correct (t(22) = 8.8, p < .0001), which reached 69% correct for the last decile of trials (t(22) = 8.2, p < .0001). However, there was an important inter-individual variability: 13 baboons out of 23 reached the criterion of 80% correct responses which was needed to pass to the next stage, whereas the remaining 10 baboons did not.

**Figure 2:**
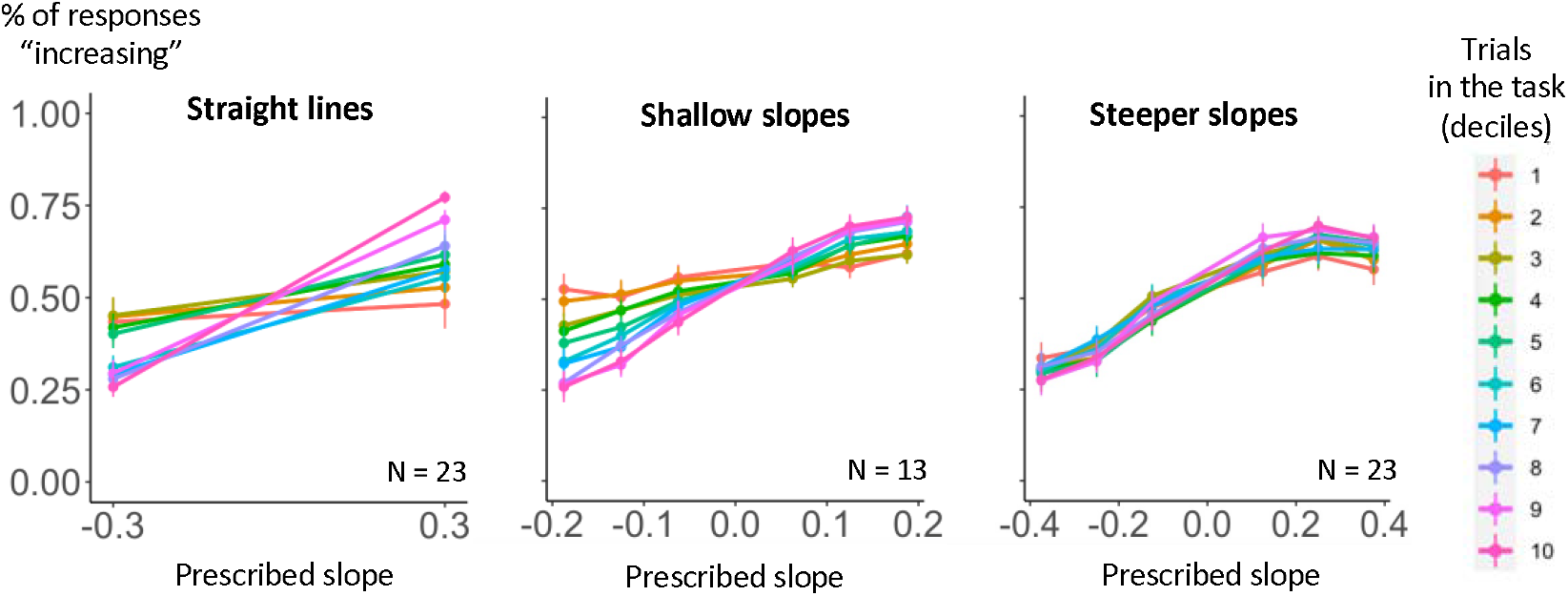
Evolution of performance on noiseless trials. Percentage of responses “increasing” with noiseless graphs in the three phases of the experiment: straight lines, noiseless lines with shallow slopes, and with steeper slopes. Responses are plotted as a function of prescribed slope. Colors indicate the evolution of time in the task, expressed as deciles of the total number of trials for a given baboon.

Once they moved to experiment 1 with noiseless lines made of 27 discrete dots, baboons poorly generalized their learning to novel stimuli: as seen in the middle panel of figure 2, their initial performance on the first decile of trial was only slightly but significantly above chance level (54% correct, t(12)= 3.8, p < .01). Over all trials, we found an accuracy of 61% correct (t(12) = 7.1, p < .0001) and an accuracy of 67% correct for the last decile of trials (t(12) = 6.8, p < .0001).

Further generalization was seen in experiment 2, when moving on to novel and steeper slopes that were presented a few weeks later, to which all baboons were exposed again, independently from their performance in experiment 1 (figure 2, right panel). The accuracy on the first decile was again significantly above chance (60% correct, t(22) = 5.9, p < .0001). The accuracy over all trials was 63% correct (t(22) = 7.7, p < .0001), and 66% correct for the last decile (t(22) = 8.4, p < .0001).

### Baboons learn to associate arbitrary shapes to increasing and decreasing trends in noisy scatterplots

We analyzed the accuracy for noisy scatterplots in experiment 1, which used the exact same stimuli previously presented to humans (Ciccione et al., 2023; Ciccione & Dehaene, 2021). We ran an ANOVA on the average percentage of “increasing” responses, with within-subject factors of slope, noise level, and number of points (except the condition with zero noise) but we found no main effects and no interactions of the factors (all p > .05). Thus, this experiment was probably too difficult for the animals, and their slightly above-chance performance did not allow to detect any determinants of behavior.

The situation was different for noisy scatterplots in experiment 2, where the graphs were steeper and less noisy. Running the same ANOVA revealed a main effect of slope (F[6, 60] = 5.88, p < .001), noise (F[3, 30] = 6.59, p = .001), and number of points (F[3, 30] = 8.36, p < .001) and an interaction of noise and number of points (F[9, 90] = 3.34, p = .001). The slope effect indicates that, as required by the task, responses depended on the actual increasing or decreasing trend of the graph. What is less easy to understand is the main effects of noise and number of points. As seen in figure S1, with 6 dots, baboons were biased to respond that the trend was increasing, but this bias reversed for larger numbers of dots. Similarly, for zero noise, baboons were, on average, consistently biased to respond that the trend was decreasing (56% of all answers), but as noise level increased, they became more biased to respond “increasing”. And finally, the interaction reflected that those biases vanished for larger noise levels. Thus, animals did not purely base their judgements on slopes, as the task required, but were biased by irrelevant parameters of noise and number of dots.

Similar findings were observed when we ran a multiple logistic regression on “increasing” answers (averaged across experimental conditions) as a function of the standardized experimental factors, now considered as numerical factors. Note that the number of points was entered (here and in the following multiple regressions) as the square root of its value minus 2, i.e., in the same way it appears in the formula of the t-value. In this regression, the actual slope and the number of points were both significant (β_actual_slope_ = .82, p < .0001; β_n_points_ = -.15, p < .05), while the noise level had no main effect (β_noise_ = -.09, p = .2). Adding interactions terms led to finding a significant interaction between the slope and the number of points (β = .17, p < .05).

To further elucidate the role of the number of points on baboons’ performance, we plotted the responses as a function of all combinations of noise and number of points (see supplementary figure S1). When there were only 6 points in the dataset, baboons behaved close to chance level. However, running the above ANOVA restricted to the 6-point conditions still showed an effect of the slope (F[6,60] = 3.56, p < .01), but no effect of noise (F[3, 30] = .28, p = .84) and no interaction of the two factors (F[18, 180] = 1.15, p = .31). Analogously, a multiple logistic regression revealed only a main effect of the slope (β_actual_slope_ = .72, p <.01). Thus, even with 6-dot graphs, baboons were able to perform the task and treat the graph as a whole. Furthermore, for the other three larger values of numbers of dots, the effect of slope became much more visible and significant (18 points: β_actual_slope_ =1.22, p < .0001; 38 points: β_actual_slope_ = 1.02, p < .0001; 66 points: β_actual_slope_ = .98, p < .0001; see figure S1).

### Baboons use the t-value in their trend judgments

For each graph, the t value combines the signed slope, the noise level and the number of points into an index of correlation strength, which was previously found to drive human behavior in the trend judgement task (Ciccione et al., 2023; Ciccione & Dehaene, 2021). We therefore plotted baboon’s “increasing” responses to noisy scatterplots as a function of the t value (for noiseless plots, the t value would be infinite and undefined). In experiment 1, logistic regression showed that performance was significantly predicted by the standardized t-value (β_t_value_ = .37, p < .001; see figure 3, top row). Accuracy was however far lower than the one observed for humans with the exact same stimuli (data from 6-year-old children, the Himba and schooled adults are presented at the top of figure 3 for comparison purposes). In experiment 2 with steeper slopes, performance was again modeled by a logistic function of t value (β_t_value_ = .82, p < .0001; see figure 3, bottom row).

**Figure 3:**
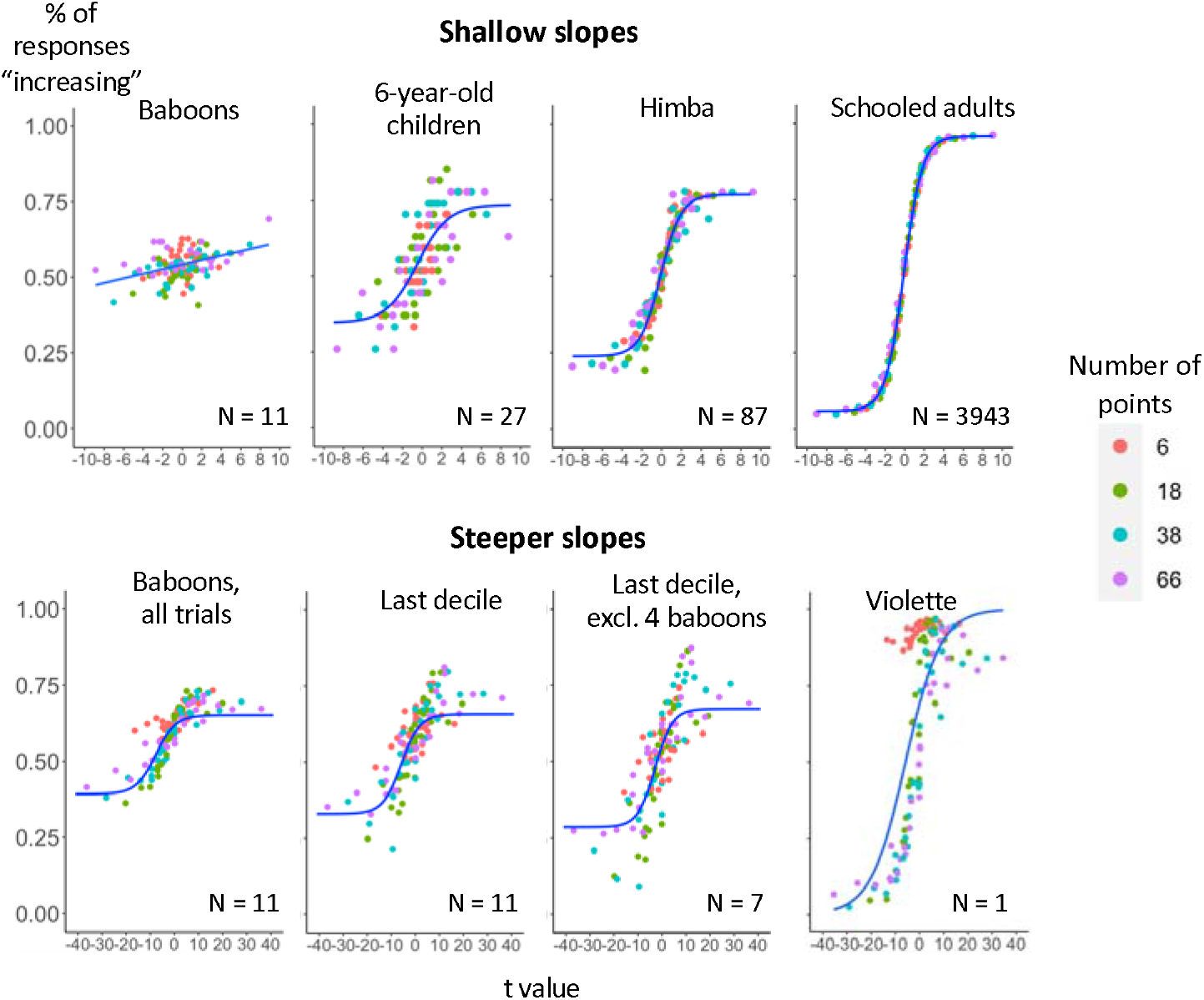
Performance with noisy graphs. Each point represents a combination of experimental factors (prescribed slope, noise, and number of points). The percentage of responses “increasing” is plotted as a function of the t-value of the scatterplot (x axis) and the number of dots (color). Top: experiment 1 (shallow slopes). Responses from all trials are plotted, together with (for comparison purposes) responses from 6-year-old children, Himba and schooled adults, as collected in a previous study which used identical stimuli (Ciccione et al., 2023). Bottom: experiment 2 (steeper slopes). Responses from all trials are plotted, together with responses from the last decile of trials, last decile excluding 4 underperforming baboons, and from Violette, a baboon who performed strikingly similarly to humans.

If the t value accurately summarizes the effects of slope, noise level and number of points, then these other variables should no longer have a significant influence on behavior once t value is entered – and indeed this was the case in human subjects (Ciccione & Dehaene, 2021).

To test this in experiment 1, we ran a multiple logistic regression model on “increasing” responses, with subjects as a random factor, now including all parameters as standardized predictors: t-value, actual slope, noise level, and number of points (again entered as 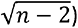. The 4 regressors were standardized (converted to z scores), so that their betas could be compared as estimates of effect size. We found only a significant effect of the t value (β_t_value_ = .49, p < .05).

We then ran the same regression for experiment 2 and we found a significant effect of t value (β_t_value_ = .32, p=.02), but also of the actual slope (β_actual_slope_ = .55, p < .0001), a close to significance effect of number of points (β_number_points_ = -.14, p = .05), but no effect of noise level (β_noise_ = .07, p = .32). This finding indicates that baboon behavior is influenced by more variables than human behavior, but it also shows that, even after controlling for the effect of other variables, the effect of the t value remains significant.

We also examined the influence of training in experiment 2. As visible in figure 3, the psychometric logistic function became steeper when we considered only the last decile of trials per subject (Figure 3, bottom row, second plot, β_t_value_ = .85, p < .0001). Indeed, this improvement in behavior was reflected in a significant interaction between t value and trial decile (β_t_value_ = .52, p < .0001; β_t_value X decile interaction_ = .12, p < .0001).

When we further excluded 4 baboons who appeared to perform at chance level (overall accuracy < 55%), the percentage of “increasing” responses became very strongly dependent on the t value (β_t_value_ = 1.14, p < .0001), with a ceiling and floor performance of, respectively, about 75% and 25% (i.e., there were only about 25% of mistakes for extremely positive or extremely negative t-values). However, even in this case, the same multiple regression as above indicated that behavior was not just influenced by the t value, but also by the actual slope and the noise level (β_t_value_ = .51; p < .01; β_slope_ = .66; p < .0001; β_noise_ = .30; p < .001).

### Inter-individual variability between baboons

When we looked at the data separately for each subject (figure S2), we found that there was a large variability between baboons, in terms of both their sensitivity at performing the task and the biases that they exhibited. By fitting a separate logistic regression for each baboon, and using the non-normalized t value as regressor, we found that the slope of the psychometric function, measuring the efficiency of their “graphicacy” (Ciccione et al., 2023), ranged from .02 for Nekke to .14 for Violette. The latter baboon’s performance (figure 3, bottom row, fourth plot) showed an almost perfect dependency on the t value (slope = .14, p < .0001). As a comparison, the median sensitivity index for 6-year-old children that we found in another study (Ciccione et al., 2023) was a slope of .12, thus lower than that of Violette.

Figure S2 shows that one major source of variability between baboons lied in the biases that they exhibited. In particular, in some baboons a larger number of points strongly favored the “increasing” response, whereas in others this effect was reversed. This bias was particularly pronounced for datasets with 6 points. However, when we ran separate multiple regressions for each subject, restricted to the stimuli with only 6 points, we found that for 6 baboons out of 11, responses were still significantly predicted by the t value. In other words, this finding suggests that on top of a strong bias, the effect of t value remained.

### Some baboons failed to perform the task because their choice stimuli were too hard to distinguish

We lastly investigated the reasons why some baboons failed to pass the first threshold of 80% correct responses with noiseless lines, and, therefore, did not perform the actual experiments with noisy scatterplots. After ruling out some trivial hypotheses, such as refusing to perform the task, inverting the response buttons, or failing to receive any reward, we looked at the pairs of shape that each baboon had to learn. As explained in the methods section, two response images out of ten were randomly assigned to each baboon. When we looked at the pairs assigned to successful baboons (i.e., those who reached the testing phase) versus unsuccessful baboons, we noticed that the two images in the first list were often similar to each other within a pair in terms of luminance and/or overall structure, whereas most images of the second list were not (see supplementary figure 3 for a comparison). This observation was confirmed by a quantitative analysis. To quantify similarity, for each image, we computed the embeddings of an artificial model (CLIP – Contrastive Language-Image Pretraining; Radford et al., 2021) and calculated the cosine similarity between the embeddings of the two images in the pair. The average similarity was significantly smaller for successful than for unsuccessful baboons (two samples t-test, t(19.2) = 3.09, p < .01). Thus, we conclude that some baboons were unsuccessful because their response stimuli were too similar to be easily distinguished. In other words, there were two sources of difficulty in the task: perceiving the ascending or descending trend of the graph, and then correctly choosing the corresponding visual stimulus – and the second was the source of failure for some animals. Conversely, for those who passed the training phase, we found no correlation between their graphicacy score (slope of the psychometric function of t value) and the discriminability of their pairs of response stimuli, suggesting that they exhibited genuine differences in the graphic perception component of the task.

## Discussion

In a series of behavioral studies, we trained baboons to learn a match-to-sample binary task in which they had to associate two arbitrary shapes, randomly appearing to either side of the screen, with the upward or downward slope of straight lines as well as noiseless and noisy scatterplots. We found that, if the animals were provided with easily distinguishable response stimuli, they could learn this task. Their performance, in the presence of noise, was predicted by the t-value of the scatterplot, the index that a statistician would use to compute the strength of the correlation in the plot.

The first aspect worth highlighting is that our experiment provides the first evidence for the ability to extract statistical information from a noisy graph in non-human animals. This skill adds to the statistical abilities already found in primates and non-primates, as reviewed in the introduction, and confirms its phylogenetically old character. It is important to underline, however, that the trend judgment task learned by our baboons is a very peculiar type of task, that needs to be distinguished from both ensemble perception and statistical learning tasks. Ensemble perception requires the extraction of the *average* value of some perceptual feature, whereas our task requires evaluating the *correlation* or covariance of two values, which is much more sophisticated. Statistical learning, on the other hand, involves the learning of regularities over time or space, following the repeated presentation of multiple stimuli which are presented serially – whereas in the current study baboons learned to extract the correlational structure of a *single* display comprising multiple items. Future studies will be needed to minimize the differences between those tasks, for instance by investigating non-human primates’ abilities to also perform classic ensemble perception tasks, such as: finding the average color hue (Webster et al., 2014), the average size (Chong & Treisman, 2003) or the average orientation (Parkes et al., 2001) of visually presented items, through both serial and parallel presentations, both with and without deviant items (e.g., Avci & Boduroglu, 2021; Ciccione et al., 2022).

Baboons’ performance with noisy scatterplots exhibited both striking similarities and differences with humans. While humans show no difficulty at extracting the overall trend of a graph comprising only 6 data points (Ciccione et al., 2023; Ciccione & Dehaene, 2021), in this condition baboons exhibited a strong bias to respond either “increasing” or “decreasing” on most trials. Crucially, however, even with 6 points the performance of many baboons was affected by the t value of the dataset, meaning that they were still performing the expected task. The difficulty with 6 points suggests that, when items are few and far from each other, baboons struggle to consider them as a single set or line. This failure is consistent with previous findings showing that baboons tend to emphasize local, item-by-item processing over the global processing of overall shape (Deruelle & Fagot, 1998; Fagot & Deruelle, 1997). Humans, on the contrary, manage to bridge across perceptual distances and to perform the task independently of the number of items. It is reasonable to attribute this difference to the fact that graphs are a cultural product and that seeing 6 dispersed points as a single graph is a culturally learned skill, or at least was an explicitly instructed task for the adults and the preschoolers that we tested. Similar considerations may apply to very noisy datasets. Baboons performed the task with a much lower accuracy when the level of noise was high (figure 3, top row), suggesting that if the points are too dispersed, they are less likely to perceived them as a single graph with a consistent trend. Future studies could investigate the limits of trend judgment in non-human primates by more finely varying the noise parameter and the number of points.

Baboons were however very similar to humans in that performance was well predicted by the t-value of the scatterplot. This finding is interesting for two reasons. First, this similar behavior, observed in both humans and baboons, suggests that the human visual system, when performing trend judgments over noisy datasets, recycles phylogenetically older brain areas involved in the recognition of the principal axis of objects (Ciccione et al., 2023). Second, it reveals that the baboons are sensitive to fine details of their statistical environment since, in order to compute the t-value, it is necessary to integrate the signed slope, the number of points and the level of noise. However, it is worth noting that, whereas for humans the responses are entirely subsumed by the t-value, this is not the case for baboons: in fact, a multiple logistic regression on their answers as a function of stimulus parameters revealed that both the t value and the actual slope of the dataset significantly affected their performance. In other words, it seems that baboons relied more on the slope than humans. This is clearly visible in supplementary figure 1: the effect of noise and number of points is in fact limited compared to the effect of the slope.

Lastly, as in humans, sensitivity in the trend judgment task varies among individuals. This finding should push animal cognition researchers to always consider inter-individual variability when studying the performance of non-human animals (Stamps et al., 2012). It would also be interesting to investigate whether such variability correlates, like in humans, with other behavioral measures, and whether or not it acts as a predictor of higher-level cognition.

## Acknowledgments

This project received funding from the European Research Council (ERC Grant to Stanislas Dehaene) and from the Agence Nationale de la Recherche ANR-16-CONV-0002 (ILCB). We thank Julie Gullstrand and Sebastien Barniaud as well as the team from the “Station de primatologie” for their help in conducting the study.

## Author contributions

Conceptualization: LC, TDB, NC, JF, SD.

Data collection: NC, JF.

Data analysis: LC

Project supervision: SD

Writing – first version: LC

Writing – revision: LC, SD.

---

**Figure S1:**
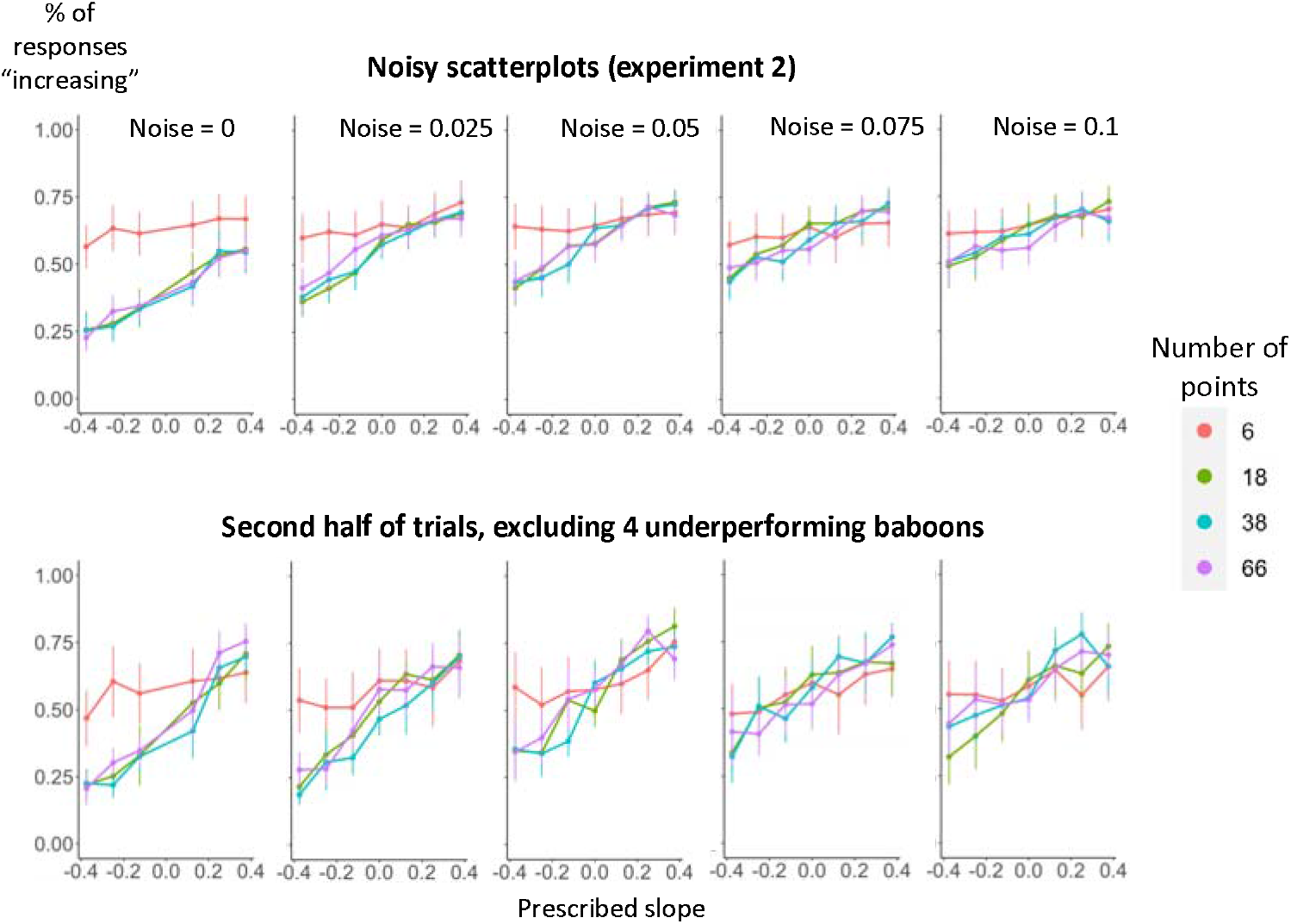
Percentage of responses “increasing” as a function of prescribed slope, noise and number of points. The first row shows the performance of all baboons from all trials. The second row shows the performance restricted to the second half of trials, only for baboons whose performance indicated that they were engaged in the task (accuracy higher than 55%).

**Figure S2:**
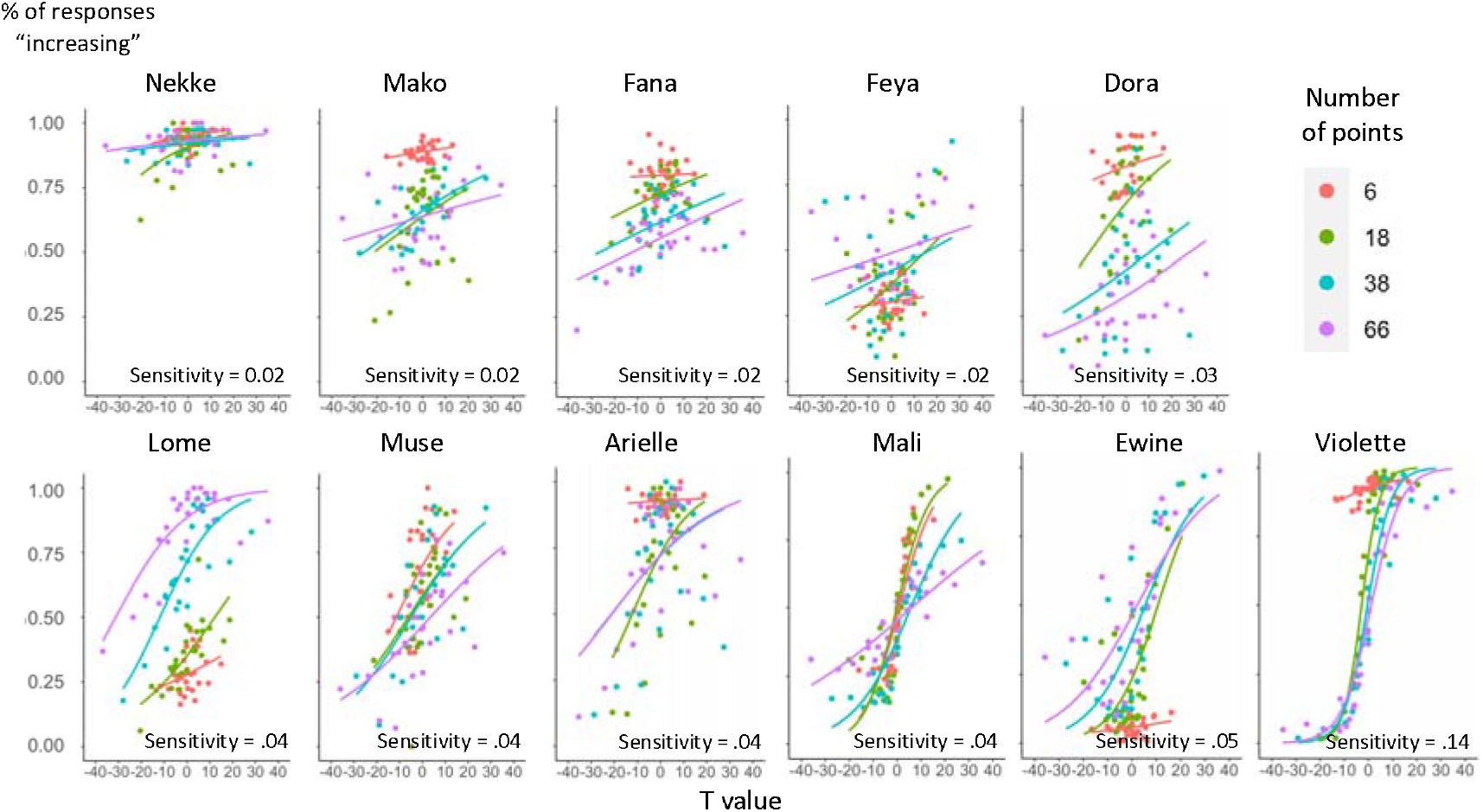
Inter-individual variability in final performance. For each subject in experiment 2 (noisy scatterplots, steeper slopes), the percentage of responses “increasing” in the second half of trials is plotted as a function of the t-value of the scatterplot (x axis) and the number of dots (color). A label indicates each subjects’ sensitivity, calculated as the slope of the sigmoid fit of given responses as a function of t value.

**Figure S3:**
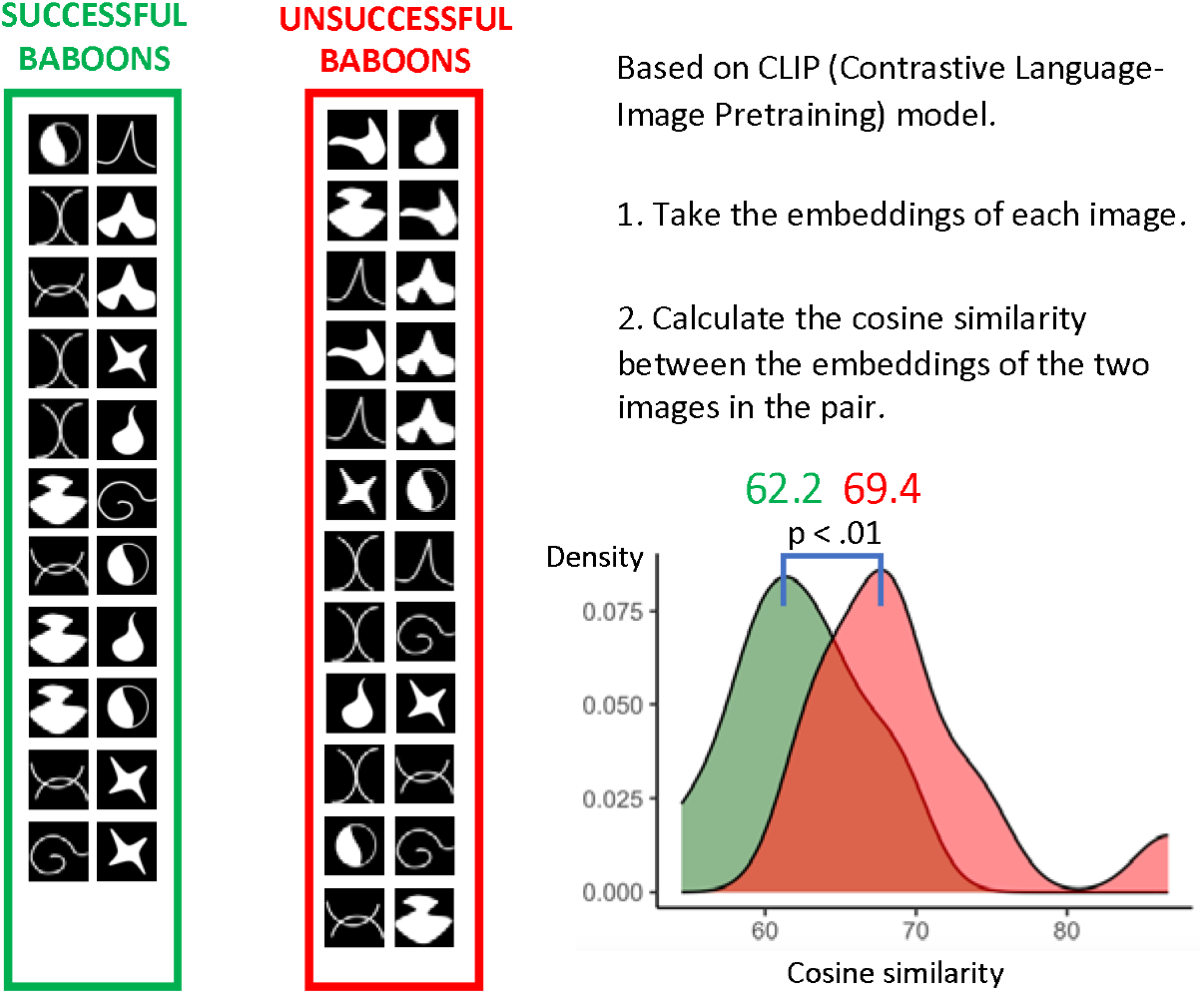
Differences in the pairs of shapes presented to each baboon. For each animal, the images show the shapes assigned to the “increasing” and “decreasing” responses (respectively left and right). The left column, in green, shows the pairs presented to baboons who reached the criterion for the task with noiseless scatterplots. The right column, in red, shows the pairs presented to baboons who failed to reach criterion. The average similarity of pairs presented to successful baboons (62.2) was significantly smaller than the average similarity of pairs presented to unsuccessful baboons (69.4). The distribution of similarities is shown at bottom right.

